# Cross-fostering affects microglia and cell death in the hippocampus of female and male degu pups

**DOI:** 10.1101/2025.08.05.668723

**Authors:** Gurprince Attlas, Mallory Duprey, Cassandra Carlson, Kaja S. Arusha, Krystle D. Boadi, Sabrina S. Ellah, Daniela Kim, Charlis Raineki, Carolyn M. Bauer, Paula Duarte-Guterman

## Abstract

Parental care is essential for social, behavioural, and neural development of offspring. Disruption of parental care, for example through parental separation, can negatively affect offspring development in rodents. While previous work has focused on maternal and paternal deprivation, the effects of cross-fostering, another form of parental-offspring instability, on brain development remain poorly understood. Such disruptions are thought to induce stress, which in turn can suppress neurogenesis and increases inflammation and apoptosis in the hippocampus, with effects that often differ between sexes. Given that degus (*Octodon degus*) are born precocial and form strong attachments with their parents early in life, this study aimed to investigate the effect of cross-fostering on hippocampal development in female and male degu pups. At postnatal day 8, degus were assigned to either control (pups remained with parents and littermates), partial cross-foster (one pup per litter was cross-fostered), or full cross-foster (the entire litter was cross-fostered) conditions. At weaning (5-weeks-old), offspring brains were collected for immunohistochemistry to examine dentate gyrus volume, density of pyknotic cells, density of immature neurons, and number and morphology of microglia. Both types of cross-fostering reduced hippocampal cell death in both sexes relative to controls, while no significant effects were observed in the dentate gyrus volume or density of immature neurons. Cross-fostering did not affect total microglia density in either sex however full cross-fostered females had fewer amoeboid microglia compared to female controls. Together, these findings indicate that cross-fostering affects hippocampal microglia and cell death in pups, potentially disrupting circuit refinement and plasticity during development, with effects that vary by cross-fostering type, sex, and hippocampal region.

## Introduction

Early life stress affects offspring development in a range of mammalian species, including humans (Gimsa et al., 2022; Kentner et al., 2010; Priebe et al., 2005; Raineki et al., 2019). In rodents, there are different paradigms which induce early life stress that include, but are not limited to, changing the parental environment via maternal separation, paternal deprivation, or cross-fostering (Aguggia et al., 2013; Boccia and Pedersen, 2001; George et al., 2010; Madison et al., 2022; Oomen et al., 2011). Any disruption to typical parent-offspring interactions can lead to alterations in offspring neurodevelopment (Champagne and Meaney, 2001). During postnatal development, there is high growth and refinement of brain circuits. At a mechanistic level, there are high rates of neurogenesis (i.e. production of new neurons) and regulated cell death (i.e., apoptosis; Hayashi et al., 2008; Loi et al., 2017; Sierra et al., 2014).

Moreover, microglia, the neuroimmune cells of the brain, play a very important role in refining circuits during early development, either by removing cells through phagocytosis or by supporting neuronal development and survival (Diaz-Aparicio et al., 2020; Sierra et al., 2014).

One of the brain regions that is most sensitive to early life stressors is the hippocampus. It is also one of the few regions where neurogenesis continues during the postnatal period (Kempermann, 2016). Past studies have shown that parental separation affects hippocampal neurogenesis, cell death (apoptosis), and microglia in the offspring, but the effects may depend on age, sex of the offspring, and type of separation (maternal/paternal, acute/chronic). For instance, acute maternal separation (24 h on postnatal day 3) reduces the numbers of immature neurons in the hippocampus of female rats at weaning, while increasing immature neurons in males (Oomen et al., 2009). In contrast, chronic maternal separation has been shown to reduce neurogenesis in male rats (Hulshof et al., 2011). Early life stressors can also affect programmed cell death. For example, maternal separation increases apoptosis in the dentate gyrus of rat pups (Yang et al., 2017). Thus, research suggests a potential link between dysregulated neurogenesis and apoptosis in the dentate gyrus due to early life stress, potentially mediated by microglial activity. Indeed, brief daily maternal separation increases both the number of and phagocytic activity of microglia in the hippocampus of juvenile male mice (Delpech et al., 2016; Johnson and Kaffman, 2018; Walker et al., 2013). In addition to removing cells through phagocytosis, microglia can also promote the production and survival of new neurons (neurogenesis) through the secretion of trophic factors (Diaz-Aparicio et al., 2020). Here we aimed to understand how these three key components––neurogenesis, cell death, and microglia––are regulated following a stressor during development.

Most studies to date have focused on the effects of maternal and paternal deprivation on offspring development, whereas the impact of other disruptions of parental-offspring interactions has received much less attention. Cross-fostering disrupts parent-offspring attachment, but unlike maternal separation or paternal deprivation, it does not rely on separation and removing pup’s access to a caring adult. Past research has shown that cross-fostering affects offspring body weight, emotional behaviour, cardiovascular and metabolic functions, and the endocrine stress response in different rodents species, including mice, degus, and rats (Arusha et al., 2024; Bartolomucci et al., 2004; Gomez-Serrano et al., 2001; Guan et al., 2023; Lu et al., 2009; Matthews et al., 2011). Degus (*Octodon degus)* are a particularly suitable model for studying the effects of cross-fostering on offspring development because they practice communal care and pups are born precocial and form strong attachments early in life with both parents and littermates (Colonnello et al., 2011; Guan et al., 2023). Importantly, the presence of siblings during cross-fostering may buffer the effects of separation from biological parents (Arusha et al., 2024). Accordingly, in the current study, we investigated the effects of cross-fostering either the entire litter (full cross-fostering) or a single pup (partial cross-fostering) on hippocampal development in female and male degus. We measured hippocampal development by assessing dentate gyrus volume, postnatal neurogenesis, apoptosis, and microglia prevalence and morphology. We hypothesized that (1) cross-fostering would impact hippocampal development by reducing neurogenesis, increasing cell death and microglial numbers, and altering the microglia morphological profile in both sexes; and (2) partial cross-fostering would be more detrimental to hippocampal development compared to cross-fostering the entire litter.

## Methods

### Study animals and breeding

Breeding pairs of captive-born *Octodon degus* and their pups were housed in the animal facility at Swarthmore College, in 51 x 25 x 33 cm acrylic terrariums containing corn cob or paper bedding allowing for visual observations. Animals were maintained under standard laboratory conditions (12:12 light/dark cycle, 23 ± 1 °C) with *ad libitum* access to water and food. Alfalfa hay was provided three times per week, and cages were cleaned weekly.

Degu females (n = 21) were paired with unrelated males (n = 21) for mating and remained together until pup weaning. All experiments were conducted in strict compliance with Institutional Animal Care and Use Committee at Swarthmore College and were carried out according to the guidelines of the Association for Assessment and Accreditation of Laboratory Animal Care.

### Experimental treatment

Pups were cross-fostered at postnatal day (PND) 8, thus allowing for sufficient time for the development of parental and social attachments within the nest (Colonnello et al., 2011; Guan et al., 2023). The date of parturition was taken into consideration while assigning the treatment groups as previously described (Arusha et al., 2024). Litters were assigned to one treatment condition: control (N=5 litters), full cross-fostering (N=4 litters), and partial cross-fostering (N=12 litters). In control litters, pups remained with biological parents and biological siblings for the duration of the experiment. In full cross-foster litters, all pups from one litter were exchanged with all pups from another such that pups were raised with unfamiliar parents but with biological siblings. In partial cross-foster litters, a single pup was exchanged with one pup of the same sex from another litter such that the pup was raised with both unfamiliar parents and unfamiliar siblings. Pups were fostered between litters of similar sizes and ages (<2 days in age difference). Between 1-2 pups per sex per group were used for subsequent analyses, with the exception of the male full cross-foster condition that included 3 male pups from the same litter (out of 4 litters). Sample sizes were decided based on previous work (Arusha et al., 2024; Guan et al., 2023) and are presented in Table 1 and each of the figure captions.

**Table 1.** Sample size for each group, and volume (mm^3^) of the dentate gyrus, hilus, and granule cell layer (GCL) in female and male degu pups in control, partial, and full cross-fostering conditions. Estimated marginal means are presented ± SEM.

|  | Sample Size |  | Dentate Gyrus |  | Hilus |  | GCL |  |
| --- | --- | --- | --- | --- | --- | --- | --- | --- |
|  | Females | Males | Females | Males | Females | Males | Females | Males |
| **Control** | 7 | 7 | 6.32±0.42 | 6.26±0.40 | 3.41±0.30 | 3.51±0.28 | 2.91±0.19 | 2.75±0.17 |
| **Partial** | 5 | 6 | 6.14±0.52 | 6.38±0.44 | 3.55±0.37 | 3.61±0.31 | 2.59±0.23 | 2.77±0.19 |
| **Full** | 6 | 9 | 6.05±0.45 | 6.81±0.39 | 3.26±0.32 | 3.89±0.27 | 2.80±0.20 | 2.91±0.17 |

### Tissue collection

Euthanasia was performed at weaning (∼5 weeks of age). Pups received 0.2 mL of undiluted Euthasol (pentobarbital sodium and phenytoin sodium solution), and brains were immediately dissected and split into the right and left hemispheres. The right hemispheres were placed on dry ice and frozen at –80°C for future analysis, whereas the left hemispheres (used in the present study) were immersion-fixed in 4% paraformaldehyde for 24h at 4°C. Following fixation, brains were then transferred to a 30% sucrose solution containing 0.1% sodium azide and stored at 4°C until sectioning. Coronal sections of 30 µm thickness were cut using a cryostat at –20°C. Hippocampal sections were collected in a seven-tube series containing an antifreeze solution (ethylene glycol, glycerol and 0.1M phosphate buffer saline with a pH of 7.4) and stored at –20°C. All three neural measurements (volume, neurogenesis, and microglia) were performed in parallel series from the same left hemisphere for all animals.

### Cresyl violet staining

Brain sections were stained with Cresyl violet to determine the effects of cross-fostering on dentate gyrus volume and to count pyknotic cells. Hippocampal tissue from one of the series was mounted onto Superfrost microscope slides. Cresyl violet staining was performed after the tissue had completely dried on the slides. The staining process started by dipping the slides into distilled water three times to remove residual salts, then submerged in 0.2% Cresyl violet acetate solution for 4 minutes. Following staining, the slides were placed in a differentiator solution, comprising 70% ethanol and seven drops of glacial acetic acid, for 30 seconds. The slides were then sequentially dehydrated in 95% ethanol (twice, for 1 minute each), followed by 100% ethanol for 1 minute. Finally, slides were cleared in xylene (twice, for 5 minutes each) and cover-slipped using Permount.

### Doublecortin immunohistochemistry

Brain sections were stained with doublecortin (DCX) to assess the effects of cross-fostering on postnatal neurogenesis. All staining procedures were conducted in multi-well plates under gentle shaking. To begin, sections were washed in 0.1M phosphate buffer saline (PBS, pH = 7.4) to remove antifreeze and then incubated in 0.6% hydrogen peroxide in PBS for 30 minutes. Sections were then rinsed with PBS and then incubated for 22h at 4°C in rabbit anti-DCX primary antibody (1:200 dilution; Cell Signalling, catalogue number: 10124618) diluted in 0.1M PBS containing 3% of normal goat serum (NGS) and 0.4% Triton-X. After incubation, tissue was rinsed in 0.1M PBS, then incubated for 18h at 4°C in biotinylated goat anti-rabbit secondary antibody (1:500 dilution; Vector laboratories) diluted in 0.1M PBS. This was followed by washes in 0.1M PBS and incubation for 4 hours in an avidin-biotin horseradish peroxidase complex solution (1:1000 dilution; ABC kit, Vector Laboratories). After rinsing in 0.1M PBS, tissue was incubated in diaminobenzidine in the presence of nickel (DAB Peroxidase Substrate Kit, Vector Laboratories) for 3 minutes. Stained sections were mounted on Superfrost microscope slides and allowed to dry for 2–3 days. Slides were then dipped in distilled water three times to remove salts, dehydrated sequentially in 70% ethanol (1 minute), 95% ethanol (1 minute), and 100% ethanol (1 minute), then cleared in xylene (twice, 5 minutes each). Finally, coverslips were applied using Permount.

### Iba-1 immunohistochemistry

Brain sections were stained for ionized calcium-binding adaptor molecule (Iba-1) to access the effects of cross-fostering on microglia. All staining procedures were conducted in multi-well plates under gentle shaking. Sections were first washed in 0.1M PBS (pH = 7.4) to remove antifreeze, followed by an incubation in 0.6% hydrogen peroxide in PBS for 30 minutes. Tissue was then rinsed with PBS and incubated for 22h at 4°C in rabbit anti-Iba-1 primary antibody (1:1000 dilution, Wako, catalogue number: 019-19741), diluted in 0.1M PBS containing 3% NGS and 0.4% Triton-X. The following day, sections were rinsed in 0.1M PBS, then incubated for ∼18h in biotinylated goat anti-rabbit secondary antibody (1:500 dilution, Vector laboratories) diluted in 0.1M PBS. This was followed by washes in 0.1M PBS and incubation for 4h in an avidin-biotin horseradish peroxidase complex solution (1:1000 dilution;

ABC kit, Vector Laboratories). Tissue was rinsed in 0.1M PBS followed by incubation in diaminobenzidine in the presence of nickel (DAB Peroxidase Substrate Kit, Vector Laboratories) for three minutes. Stained sections were mounted on Superfrost microscope slides and allowed to dry for 2–3 days. Slides were dipped in distilled water three times to remove salts, dehydrated sequentially in 70% ethanol (1 minute), 95% ethanol (1 minute), and 100% ethanol (1 minute), then cleared in xylene (twice, 5 minutes each). Coverslips were applied using Permount.

### Microscopy, Imaging, and Quantification

All analyses were performed by a researcher blinded to experimental conditions.

Brightfield images of the entire left hippocampus stained with Cresyl violet were taken using a standard light microscope (Olympus BX53) equipped with a camera (ORCA-spark, Hamamatsu) using the 4X objective. The granule cell layer (GCL) and hilus were traced separately using ImageJ to obtain areas in µm^2^. Volumes of GCL and hilus were calculated using the Cavalieri’s principle: the measured areas were multiplied by the section thickness (30 µm), by 7 (reflecting the 7 sections collect for the hippocampus), and finally by 2 (to present the volume for both hemispheres). The dentate gyrus volume was calculated by adding the volume of the GCL and the hilus. For the pyknotic cell count, two dorsal and two ventral sections of the dentate gyrus were selected for each animal. Ventral sections were defined as those located between interaural-incisor coordinates 2.32 and –0.32 of the degu brain atlas (Kumazawa-Manita et al., 2018). We counted the number of pyknotic cells in the GCL using a 100X oil immersion objective. During counting, the fine focus knob was used to scan through the entire thickness of each section to ensure no cells were missed.

For the analysis of immature neurons, pups exhibited a very high expression of DCX, typical for this developmental period, but making it challenging to reliably delineate and count individual immature neurons (Figure 2A and B). Therefore, we used optical density as an index of DCX expression. In addition, although morphological classification of DCX+ cells can provide insight into neuronal maturation, the high expression of DCX also prevented us from reliably scoring individual cell processes. Brightfield images were taken from two dorsal and two ventral sections using an Olympus BX53 microscope with a 4X objective. Optical density (OD) of DCX labeling was quantified using ImageJ. The GCL was traced in each image, and grey values were obtained. The Rodbard equation was employed to calculate the OD for both the GCL and background region (corpus callosum). For each section, background OD was subtracted from the GCL OD. The average corrected OD for both image densities was used in statistical analysis.

**Figure 1.**
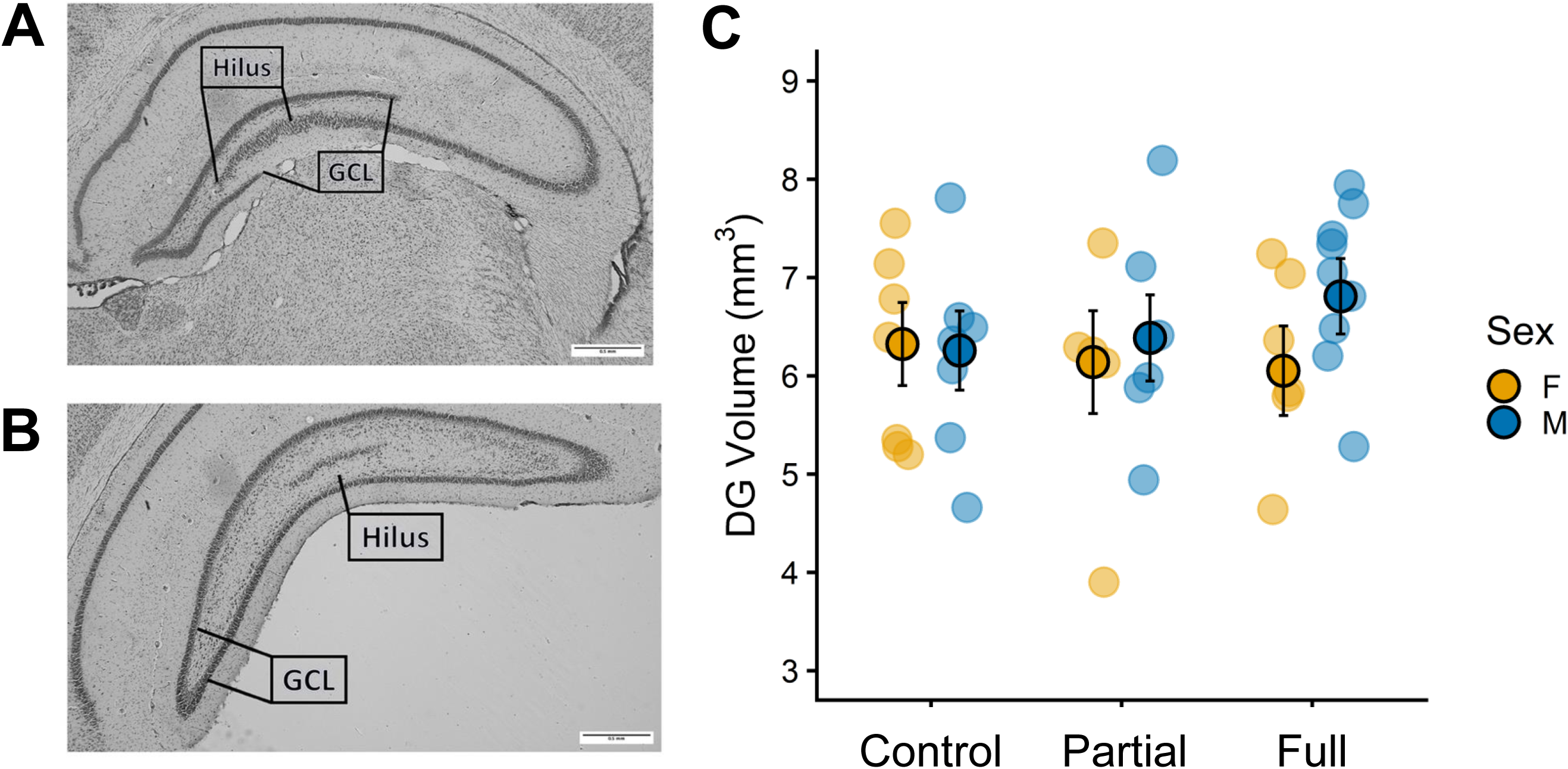
Representative image of the dorsal (A) and ventral (B) hippocampus in degu pups. The granule cell layer (GCL) and hilus are marked. Dentate gyrus volume (C) was not affected by partial or full cross-fostering in male and female pups. Individual data points are shown with the estimated marginal means (±SEM). Sample sizes were the following: control n=7 females, n=7 males; partial cross-foster n=5 females, n=6 males; full cross-foster n=6 females, n=9 males.

**Figure 2.**
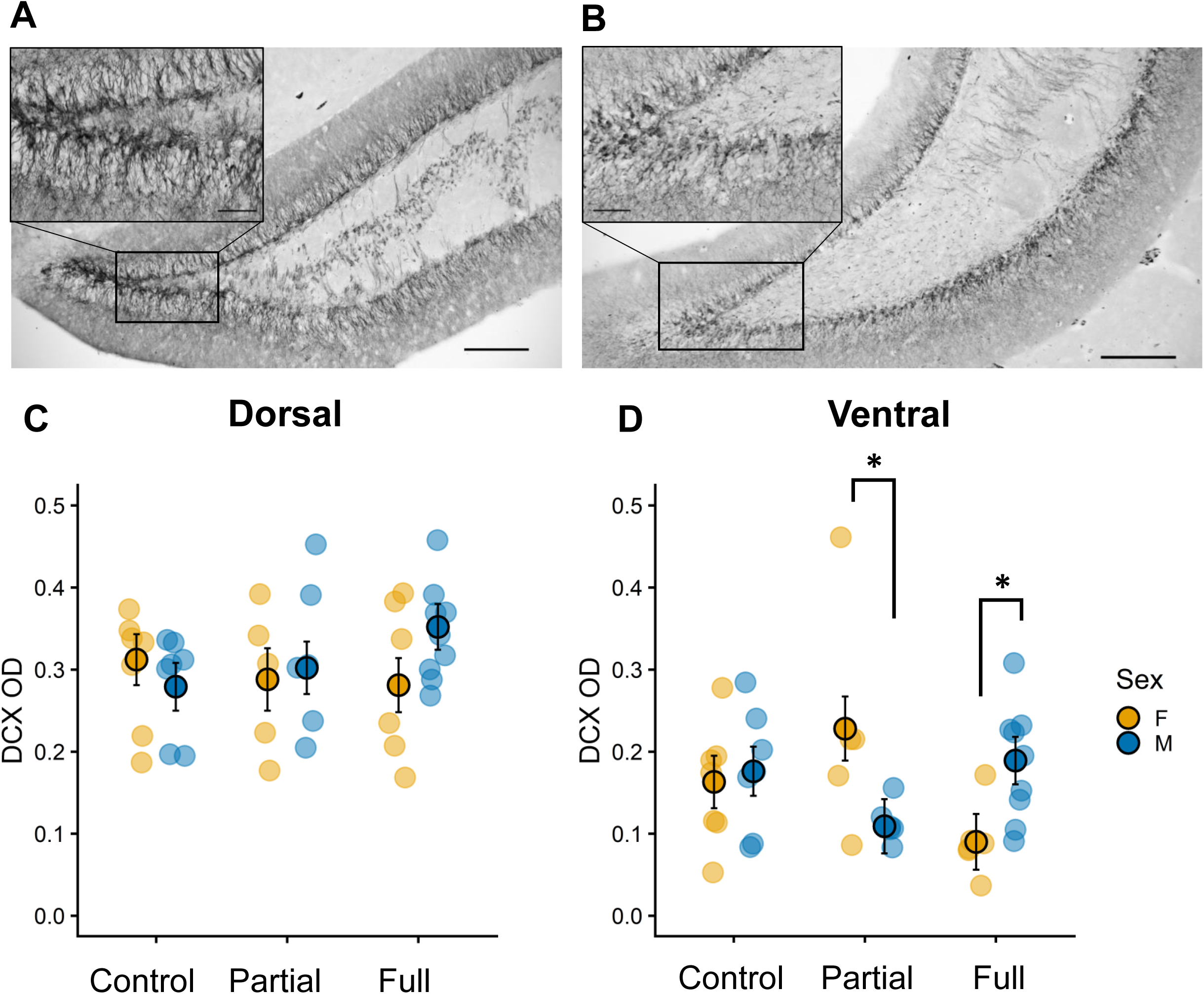
A) Representative image of DCX expression in the dorsal (A) and ventral (B) hippocampus of degu pups. Scale bars equal to 200 µm or 50 µm. C) DCX optical density (OD) in the dorsal (D) and ventral (C) regions in female and male degus. Cross-fostering did not affect DCX OD in either region in both sexes. Individual data points are shown with the estimated marginal means (±SEM). Sample sizes were the following: control n=7 females, n=7 males; partial cross-foster n=5 females, n=6 males; full cross-foster n=6 females, n=9 males.

For microglia, brightfield images were taken using an Olympus BX53 microscope with a 20X objective. Two images from the dorsal and two from the ventral dentate gyrus per animal were taken. We used a standardized region of interest (490.4 x 368.5 µm; see placement in Figure 3A), which included the GCL and hilus regions. We counted the total number of microglia present and categorized them into three morphological types—ameboid, intermediate, and ramified—using ImageJ, as we have done in the past (Duarte-Guterman et al., 2023).

**Figure 3.**
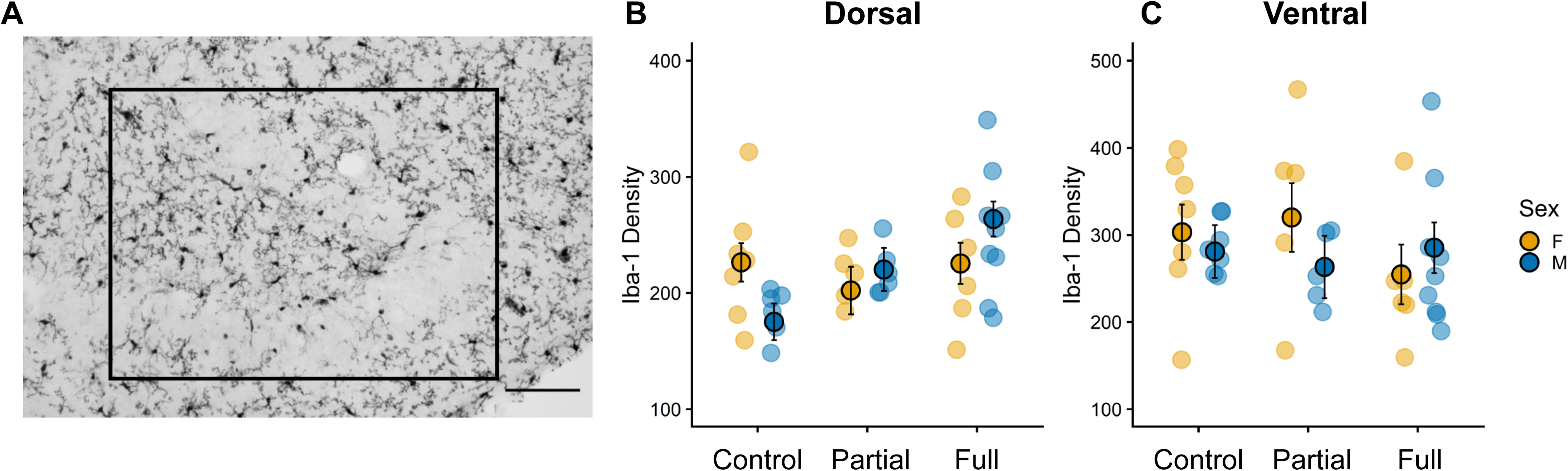
(A) A depiction of Iba-1 expression within the region of interest (black rectangle box) in the dentate gyrus of the hippocampus. Total number of microglia in the dorsal (B) and ventral (C) hippocampus. There were no significant differences across cross-fostering groups in either region. Individual data points are shown with the estimated marginal means (±SEM). Sample sizes were the following: control n=7 females, n=7 males; partial cross-foster n=5 females, n=6 males (one male sample was damaged and did not have dorsal sections); full cross-foster n=6 females, n=9 males.

Ramified microglia were defined by a small cell body with at least three processes, each at least twice the length of the cell body diameter. Intermediate microglia were defined by cell bodies with 1–2 processes, each no longer than twice the diameter of the cell body. Ameboid microglia were defined as a dark cell body with very small or no visible processes. All microglia were fit into the best or closest category by an experimenter that was blinded to the treatment groups. The number of microglia was added to get the total number of microglia per individual and normalised to region of interest to obtain a density measure. For one partial cross-fostered male, due to tissue loss during immunostaining, we did not have dorsal sections for Iba-1 analysis. One control female brain did not have two ventral section images due to missing tissue sections, therefore, data of only one section were utilized in the analysis.

### Statistical Analysis

All statistics were performed in IBM SPSS, Version 30.0.0.0 and data visualization was performed in R (R Core Team, 2021) using the ggplot2 package (Wickham, 2016). ANOVA assumptions of normality and homogeneity of the variance were checked with visual inspections of QQ plots and residual plots, and Kolmogorov-Smirnov and Levene’s tests, respectively. The volumes of the dentate gyrus, hilus, and GCL were analysed using analysis of covariance (ANCOVAs) with treatment (control, partial, and full cross-foster) and sex (female, male) as between-subject factors, and litter size and number of females as covariates. For DCX OD, microglia density, microglia morphology, and pyknotic cell density, data were analysed using repeated measures ANOVAs, with treatment and sex as between subject factors, region (dorsal and ventral) as a within-subject factor, and litter size and number of females as covariates. Including litter size and sex ratio as covariates ensured that treatment-related effects were not confounded by natural variation in litter composition. Including these covariates in the analysis did not affect the main patterns of the results. For all tests, we present the estimated marginal means and the raw data. Post hoc comparisons for the repeated-measures ANOVA were conducted using the Sidak correction. For the repeated measures ANOVAs, if the assumption of sphericity was violated, the degrees of freedom were corrected using the Greenhouse-Geisser estimate of sphericity. Pearson r correlations were performed between cell types, separately for females and males and the different treatment groups. P-values < 0.05 were considered statistically significant.

## Results

### Cross-fostering did not alter dentate gyrus volume or DCX optical density

Cross-fostering did not affect the volume of the dentate gyrus in female and male pups, (all p’s > 0.526) (Figure 1). Similarly, cross-fostering did not affect the volume of the hilus (all p’s > 0.292) or the GCL (all p’s > 0.595; Table 1). We did not find any sex differences for any of the volumes (dentate gyrus, hilus, or GCL) (all p’s > 0.292; Table 1).

To examine the effects of cross-fostering on postnatal neurogenesis, we measured the OD of DCX in the dorsal and ventral regions of the hippocampus. We found a significant interaction between treatment, sex, and region (F (2,32) = 4.212, p = 0.024; η^2^ = 0.208). Post hoc analysis indicated that partial cross-fostered females had a trend for significantly higher DCX OD than full cross-fostered females in the ventral hippocampus (p=0.059; Figure 2). However, this trend was largely driven by a single outlier in the partial cross-fostered group; after winsorising this value to the nearest non-extreme value, the contrast was no longer significant (p = 0.458). There were no significant differences between partial cross-fostered females and control females (p = 0.168), or between full cross-fostered females and control females (p = 0.157) in the ventral hippocampus. There were no differences between males across treatment conditions in DCX OD in the ventral hippocampus (all p’s > 0.072). In the dorsal hippocampus, cross-fostering did not affect DCX OD in either females or males (all p’s > 0.101).

Furthermore, we detected sex differences in DCX OD in the partial cross-fostered and full cross-fostered pups. In the ventral hippocampus, partial cross-fostered females had higher DCX OD than males (p = 0.032) although this sex difference disappears when the partial cross fostered female outlier is replaced with the nearest non-extreme value. In contrast, full cross-fostered females had lower OD compared to males (p = 0.023). In controls, no sex differences in DCX OD were found in either the dorsal or ventral hippocampus (all p’s > 0.429).

### Cross-fostering altered microglia counts and morphology in the dentate gyrus of degu pups

Cross-fostering did not affect the density of Iba-1-positive cells in the dentate gyrus of degu pups when taking into account litter size and sex ratio in the analysis (all p’s > 0.149, including the effect of covariates; Figure 3). Without these covariates, there was a significant treatment by region interaction: (F (2,33) = 3.824, p = 0.033), however post-hoc analyses did not reveal any significant differences between treatments.

While total Iba-1 density was not affected by cross-fostering, we analysed the density of the three morphological states of Iba-1 expressing cells. Cross-fostering also affected the density of ameboid Iba-1-positive cells with the effect being sex-specific but across both dorsal and ventral regions (treatment by sex interaction: F (2,31) = 4.531, p = 0.019; η^2^ = 0.226, Figure 4). Full cross-fostered female pups had fewer ameboid microglia compared to female controls (p = 0.026). Partial cross-fostering did not affect the density of ameboid Iba-1 positive cells in female pups compared to control or full cross-fostering conditions (all p’s > 0.449). No significant differences were found in the density of ameboid microglia in males (all p’s > 0.733). No significant differences were observed in the density of ramified or intermediate microglia across cross-fostering conditions (all p’s > 0.089).

**Figure 4.**
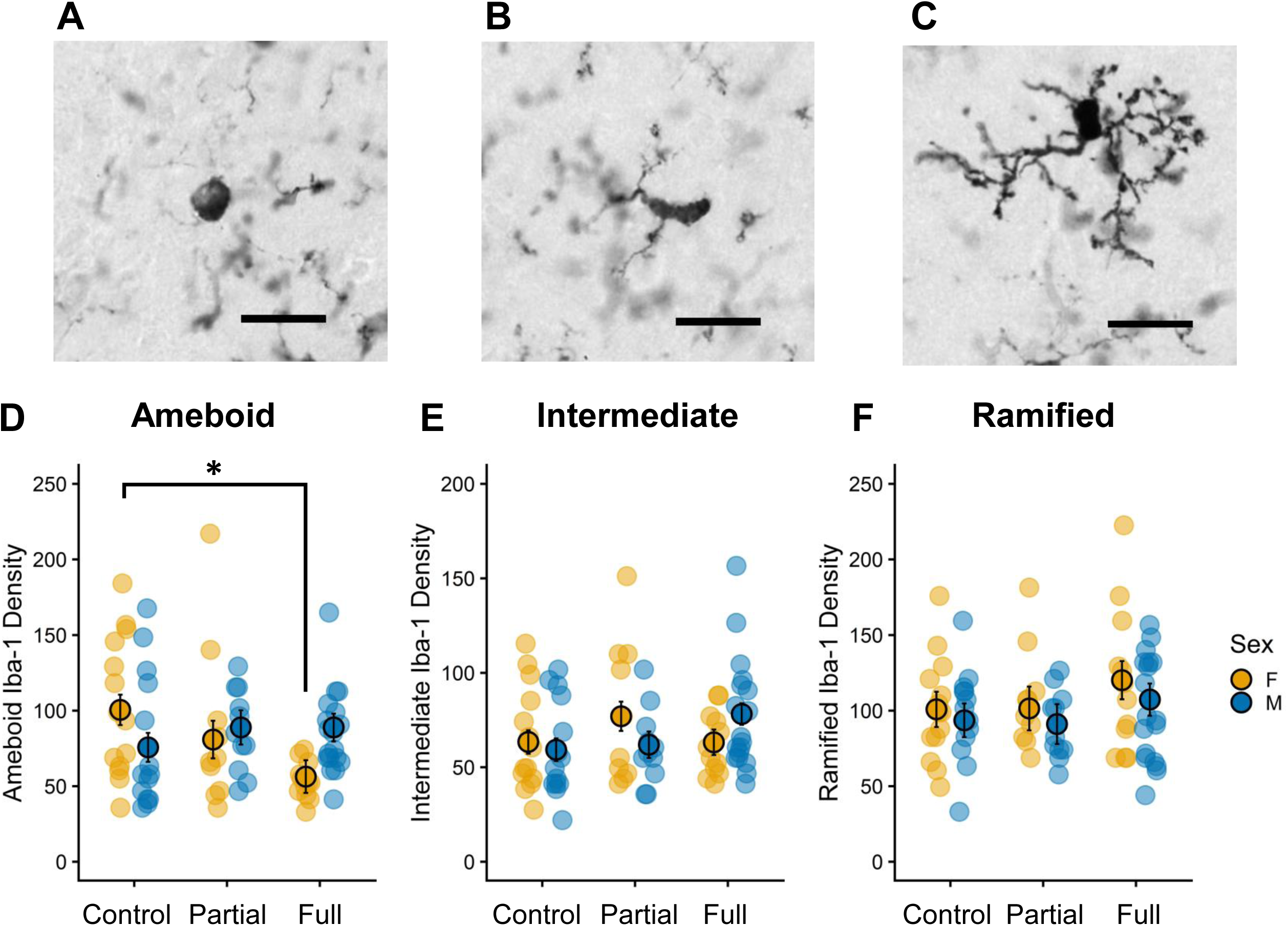
Representative images of microglia morphology showing (A) ameboid, (B) intermediate, and (C) ramified microglia (scale bar = 20 µm). (D) Full cross-fostered female pups had a lower ameboid microglia density than control (* p < 0.05). Cross-fostering did not affect the density of intermediate (E) or ramified microglia (F). Individual data points are shown with the estimated marginal means (±SEM). Sample sizes were the following: control n=7 females, n=7 males; partial cross-foster n=5 females, n=6 males (one male sample was damaged and did not have dorsal sections); full cross-foster n=6 females, n=9 males.

### Cross-fostering differentially altered pyknotic cell density in dorsal and ventral hippocampus regions

Cross-fostering affected pyknotic cell density differently in the dorsal and ventral hippocampus (treatment by region interaction: F (2,32) = 5.058, p = 0.012, η^2^ = 0.240, Figure 5). In the dorsal and ventral hippocampus, full cross-fostered pups had fewer pyknotic cells compared to controls (p < 0.001 and p = 0.025, respectively). Partial cross-fostered pups also had fewer pyknotic cells compared to controls but only in the dorsal hippocampus (p = 0.002) No sex differences were found in pyknotic cell density in either dorsal or ventral regions (all p’s > 0.127).

**Figure 5.**
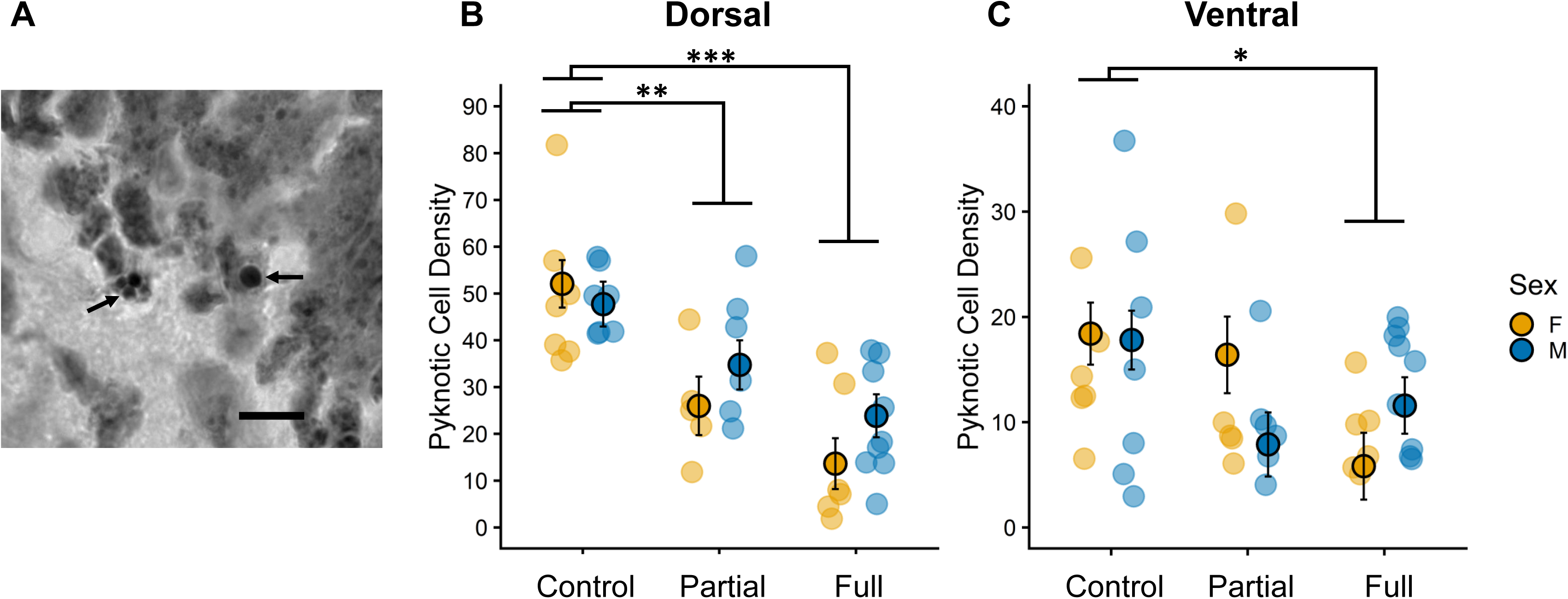
(A) Representative image of pyknotic cells (arrows) in the granule cell layer in degu pups (scale bar = 20 µm). Pyknotic cell density was affected by cross-fostering in the dorsal (B) and ventral hippocampus (C). In the dorsal region, full cross-fostered pups had the lowest density of pyknotic cells, followed by partial cross-fostered pups when compared to controls (** p < 0.01, *** p < 0.001). In the ventral region, full cross-fostered pups had a lower density of pyknotic cells compared to controls (* p<0.05). Individual data points are shown with the estimated marginal means (±SEM). Sample sizes were the following: control n=7 females, n=7 males; partial cross-foster n=5 females, n=6 males; full cross-foster n=6 females, n=9 males.

### Correlations between cell types

We examined correlations between microglia density and morphology, DCX OD, and pyknotic cell density within each cross-fostering condition, separately for males and females and hippocampal region. In the dorsal region, we found density of ramified microglia was significantly correlated with DCX OD in partial cross-fostered males only (Pearson r = 0.914, p = 0.03). No other significant associations between these endpoints in either sex were found in the dorsal region (all p’s > 0.10). In the ventral region, intermediate microglia density was correlated with DCX OD in partial cross-fostered females only (Pearson r = 0.948, p = 0.014). There were no other significant correlations in the ventral region (p’s > 0.08).

## Discussion

The purpose of this study was to examine the effects of cross-fostering on neural development in female and male degu pups. Degus are born precocial and form strong attachments early in life, making them a great model for studying the effects of early-life adversity and social bonding during development (Arusha et al., 2024; Guan et al., 2023). We found that cross-fostering affected microglia morphology and pyknotic cell density, but not dentate gyrus volume or neurogenesis. Full cross-fostering was associated with a reduction in ameboid microglia in female pups in both dorsal and ventral regions. Full cross-fostered pups, regardless of sex, had a lower density of pyknotic cells compared to controls in the dorsal and ventral regions. However, partial cross-fostering reduced pyknotic cell density in both sexes but only in the dorsal region. Overall, this study demonstrates that cross-fostering affects hippocampal microglia and cell death in a type of cross-fostering-, sex-, and region-specific manner in degu pups.

### Cross-fostering did not affect dentate gyrus volume in pups

Cross-fostering did not impact the volume of the dentate gyrus in degu pups. Previous work in other rodents such as rats, mice, and California mice has been mixed. Some studies find reductions in the dentate gyrus volume of pups exposed to early life stressors such as paternal deprivation (Madison et al., 2022), chronic unpredictable stress (Herpfer et al., 2012), and scarcity-adversity model using low bedding (Raineki et al., 2019). However, other studies find no effect after exposure to chronic unpredictable stress (Huot et al., 2002) or maternal separation (Loi et al., 2017). Overall, our findings highlight the importance of considering both the type of stressor and the species when evaluating the effects of early life stressors on hippocampus development. Notably, in the current experiment, maternal and paternal licking and grooming were not affected by cross-fostering in degus (partial or full cross-fostering), although cross-fostered mothers spent more time off the nest compared to control and partial cross-foster groups (Bauer et al., in prep). It is possible that having a relatively more stable parental care environment (e.g., compared to separation paradigms) may help protect against the effects of cross-fostering on dentate gyrus volume in pups.

Interestingly, we did not find sex differences in dentate gyrus volume in degu pups. This is in contrast with findings in mice, where males have larger hippocampal volume compared to females between PND 10 and PND 17, and again at PND 65 (Qiu et al., 2018). To our knowledge, our study is the first one to examine hippocampal volume in degu pups. Further research is required in both juvenile and adult degus to understand sex differences in hippocampal development.

### Cross-fostering does not affect neurogenesis in pups

We examined the effects of cross-fostering on postnatal neurogenesis using DCX OD as a marker of immature neurons. We found that cross-fostering did not affect DCX OD compared to controls. Previous work has found that maternal separation leads to a reduction in the number of immature neurons at weaning in female, but not male rats (Oomen et al., 2009). In biparental species, paternal deprivation either has no effect on neurogenesis (cell survival) in adult California mice (Madison et al., 2022), or decreases cell survival in adult females only (Glasper et al., 2018). In mandarin voles, paternal deprivation is associated with a reduction in immature neurons in adulthood compared to biparentally raised pups (He et al., 2018). Overall, these findings suggest that changes in neurogenesis following parental separation are not always observed in all studies and results can vary across sexes and potentially with age. Our observations were limited to a single developmental time point and one neurogenesis marker using optical density. It remains possible that cross-fostering may affect the number of immature neurons, without affecting the density of DCX, or that maturation and survival of new neurons are affected, or that changes are manifested later in adolescence or adulthood.

### Cross-fostering alters pyknotic cell density in pups

The current study revealed noteworthy alterations in pyknotic cell density in the dentate gyrus depending on the type of cross-fostering and region. We found that full cross-fostered pups, regardless of sex, exhibited fewer pyknotic cells compared to controls in both regions of the hippocampus, while partial cross-fostered pups had reduced pyknotic cell density in the dorsal region only. In contrast, previous studies in rats have shown that maternal separation can increase cell death, although the effects depend on timing of exposure. For example, maternal separation increases hippocampal cell death after an acute 24-hour separation at PND12, but not at PND20 (Zhang et al., 2002). Increases in hippocampal cell death have also been reported following chronic maternal separation in rats (Baek et al., 2012). These discrepancies between previous work and the current study may reflect species differences or differences in the type and severity of the early life stressor, as discussed earlier. Apoptosis is a crucial developmental process that contributes to the formation and refinement of neural circuits, and alterations in this process can influence neuron number and brain function (Forger, 2009; Majcher-Maślanka et al., 2019). Thus, the observed reduction in pyknotic cell density in cross-fostered pups could indicate a decreased level of apoptosis and insufficient cell pruning, potentially resulting in overcrowding of neurons and altered neural circuitry.

Partial cross-fostering affected the dorsal and ventral regions differently in pups. The dorsal hippocampus is primarily involved in spatial learning and other cognitive aspects of memory, whereas the ventral hippocampus is associated with stress and anxiety regulation (Bannerman et al., 2004; Fanselow and Dong, 2010; Kheirbek and Hen, 2011). Further research is needed to determine whether cross-fostering-related changes in cell death during development translate into changes in cognition, mood or other hippocampal-related behaviours in degus.

### Cross-fostering altered microglia numbers and morphology in pups

We found that cross-fostering altered the morphology of microglia in the hippocampus but not overall microglia density. Microglia play a crucial role in the central nervous system, contributing to synaptic remodeling through pruning, as well as supporting neuronal survival via the release of neurotrophic factors (Cornell et al., 2022; Sierra et al., 2014). Although we did not observe changes in overall microglia numbers, previous work has shown that an adverse early life environment due to a combination of minimal bedding, frequent cage changes, and a Western diet increases microglial numbers in the mouse hippocampus at weaning (Cohen et al., 2016). Similarly, maternal separation in rats (Banqueri et al., 2019) and paternal deprivation in California mice (Madison et al., 2022) have also been associated with increased microglia in the offspring hippocampus.

Instead, in the current study, we found that cross-fostering affected the morphology of microglia in a sex-dependent manner. Full cross-fostered female pups had fewer ameboid microglia compared to control females regardless of hippocampal region. Ameboid microglia are typically associated with phagocytosis, synaptic pruning, and the secretion of pro-inflammatory cytokines and are characterized by a rounded morphology lacking the long processes often observed in ramified microglia (Burke et al., 2016; Colonna and Butovsky, 2017; Franco and Fernández-Suárez, 2015; Paolicelli et al., 2022). The current literature agrees that microglia vary from ameboid to ramified states, but each morphology does not have exclusive functions. The ability to transition between different stages allows microglia to maintain a balanced immune response (Paolicelli et al., 2022). One possible interpretation of our results is that the reduction in ameboid microglia in full cross-fostered female pups may reflect lower levels of phagocytosis, potentially contributing to the reduced pyknotic cell density observed. However, this may not be the only contributing factor because cross-fostering also decreased pyknotic cell density in males with no changes in microglia morphology. In addition, there were no significant correlations between microglia (numbers and morphology) and pyknotic cell density in our samples, although the results may be limited by the small sample size. Overall, while cross-fostering clearly affected microglial morphology in the female hippocampus, whether these changes directly lead to apoptosis remain to be elucidated.

### Brain measures depend on type of cross-fostering

Our results indicate that full cross-fostered pups showed greater effects on cell death and microglia morphology, and to some extent, neurogenesis compared to partial cross-fostered pups. These findings were contrary to our hypothesis and previous work demonstrating that presence of a familiar sibling (when the entire litter is transferred) reduces the effect of cross-fostering on circulating cortisol levels and weight gain, compared to pups that were cross-fostered on their own (Arusha et al., 2024). In adult rats, the effects of corticosterone on hippocampal neurogenesis vary depending on the dose, with studies showing increases, decreases, or no effect (reviewed in Duarte-Guterman et al., 2017). It is possible that a lower-stress condition (such as full cross-fostering based on cortisol levels and weight gain; Arusha et al., 2024) may paradoxically produce greater effects on the hippocampal development compared to a moderate stressor (such as partial cross-fostering). Further work is needed to investigate the impact of stress hormones on hippocampal development in degus.

## Conclusion

Because degus are precocial and fathers, in addition to mothers, may also contribute to offspring care, studies examining cross-fostering in this species may provide a more translationally relevant model for understanding how early-life caregiving environments, such as those experienced in adoption and foster care, shape developmental outcomes. In this context, our results demonstrated that cross-fostering did not affect volume or postnatal neurogenesis in the hippocampus. However, cross-fostering affected pyknotic cell density and microglia morphology, in a type of cross-fostering-, region- and sex-specific manner. Our findings complement earlier work with the same cross-foster degu model to examine its effects on HPA axis regulation (Guan et al., 2023; Arusha et al., 2024) and they further advance our understanding of how this early life stressor influences brain development. Finally, the results also highlight the importance of including both sexes and diverse forms of early-life stressors, such as cross-fostering paradigms. Further work is needed to determine the long-term effects of cross-fostering on brain function and potential behavioural consequences.

## Acknowledgments

This research was supported by the Canada Research Chair Program to PDG and Swarthmore College to CMB. GA was funded by Ontario Graduate Scholarship and Faculty of Social Sciences at Brock University. SE was funded by Women’s Science Research Fellowship and KB, DK, and KA were funded by Swarthmore College Summer Opportunities and Research Fellowships. We thank the animal care staff at Swarthmore College for technical assistance.

